# Unveiling the Impact of Vernalization on Seed Oil Content and Fatty Acid Composition in Rapeseed Through Simulated Shorter Winters

**DOI:** 10.1101/2024.01.04.574184

**Authors:** İrem Çağlı, Büşra Elif Kıvrak, Osman Altunbaş, Çağla Sönmez

## Abstract

Climate change is leading to warmer winters world-wide with an increasing number of extreme events every year. Plants are majorly impacted by the escalating effects of global warming. In this study, we set up an experimental model to simulate warmer and shorter winters under laboratory conditions. Winter and spring varieties of rapeseed (*Brassica napus* L.) were subjected to diverse vernalization scenarios including three and four weeks-long vernalization as well as vernalization interruptions by one week-long devernalization at warm temperatures. The aim of the study was to assess the effects of the ‘vernalization models’ on *BnaFLC* (*BnaFLCA02*, *BnaFLCA10* and *BnaFLCC02*) expression, some yield related traits, a set of genes involved in fatty acid synthesis and seed oil content and fatty acid composition. A notable difference in vernalization responsiveness was observed in *BnaFLCA02*, *BnaFLCA10*, and *BnaFLCC02* between the late-flowering winter variety, Darmor, the early-flowering winter variety Bristol and the spring variety, Helios, after a three-week vernalization period. Our findings unveil a robust correlation between vernalization and seed oil content, as well as fatty acid composition in rapeseed. While the expression levels of fatty acid synthesis-related genes, including *BnaFAD2*, *BnaFAD5*, *BnaFATB*, *BnaMCOA* (*AAE13*), and *BnaWD40*, exhibited significant changes under cold conditions in leaves, the expression levels of the same genes in developing seeds did not exhibit a strong correlation with vernalization, flowering time, or oil and fatty acid contents in seeds. Our results suggest that vernalization plays a role in seed oil biosynthesis beyond its impact on flowering time.

---

1. **Introduction**

Rapeseed (*Brassica napus* L.) is an economically important oilseed crop valued for its high seed oil content ranging from 35% to 50%. The adaptability of both winter and spring rapeseed varieties to diverse environmental conditions significantly contributes to its widespread cultivation (Marjanovic-Jeromela et al., 2019).

The winter varieties need a prolonged cold period, known as vernalization, to transition from the vegetative to the generative growth phase in a timely manner. This vernalization process allows plants to monitor environmental signals, ensuring optimal flowering and subsequent seed development under favorable conditions for reproductive success. Extensive molecular analyses conducted on *Arabidopsis thaliana* have revealed the intricate nature of vernalization responses in plants. Key determinants of vernalization-induced flowering time in Arabidopsis include the *Flowering Locus C* (*FLC*) and *FRIGIDA* (*FRI*) genes, which exhibit antagonistic effects on the vernalization response in plants (Napp-Zinn, 1979; Koornneef et al., 1991; Clarke & Dean, 1994; Koornneef et al., 1994; Michaels & Amasino, 1999; Shindo et al., 2005).

The specific role of FRI in vernalization is to directly bind to the *FLC* mRNA, keeping *FLC* gene expression high (Michaels & Amasino, 2001; Geraldo et al., 2009). FLC acts as a transcriptional repressor and regulates the expression levels of many genes associated with flowering time such as *FLOWERING LOCUS T* (*FT*) and *SUPPRESSOR OF CO* (*SOC1*), which are involved in floral transition (Whittaker & Dean, 2017). Consequently, *FLC* functions to inhibit flowering. During vernalization, *FLC* gene expression gradually decreases from the second or third week onwards, and this reduction is mediated by an epigenetic mechanism that involves the Polycomb Repressive Complex 2 (PRC2) and nucleation (Bastow et al., 2004; De Lucia et al., 2008; Swiezewski et al., 2009). Additionally, a chromatin loop formed by the 5’ and 3’ regions of the *FLC* gene is disrupted during the early stages of vernalization (Crevillen et al., 2013).

*Brassica napus* (2n = 4x = 38; genome AACC) is an allotetraploid belonging to the Brassicaceae family like *A. thaliana* (Chalhaub et al., 2014). Many genes related to vernalization and flowering time are conserved between Arabidopsis and *B. napus* (Schiessl et al., 2014). Unlike the single *FLC* copy in *A. thaliana*, *B. napus* genome has nine copies of the *FLC* gene (Zou et al., 2012). Specifically, *BnaFLCA02*, *BnaFLCC02*, and *BnaFLCA10* exhibit differential expression in vernalization-dependent versus vernalization-independent varieties and are downregulated by vernalization (Schiessl et al., 2019). While *BnaFLCA02* and *BnaFLCC02* expression delay flowering in spring varieties, they do not prevent it (Tudor et al., 2020). Genome-wide association study (GWAS) of 188 different Brassica accessions revealed that natural variation in flowering time is primarily determined by *BnaFLCA10* or *BnaFLCA3a* accounting for 15% and *BnaFLCC02* accounting for 23% of the variance (Raman et al., 2016). QTL-seq analysis further identified *BnaFLCA02* accounting for flowering time variation in winter rapeseed cultivars with distinct vernalization requirements (Tudor et al., 2020). Although various studies highlighted the relative contributions of each *FLC* gene to the vernalization response and the flowering time, a modeling approach revelead that the total expression levels of nine *FLC* paralogs collectively contribute to the vernalization response in different rapeseed varieties (Calderwood et al., 2020).

Climate change poses a significant threat to plant production, leading to considerable crop losses. (Lesk et al., 2016). Nationwide yield data from the United Kingdom demonstrated a strong correlation between a 1 °C increase in early winter temperature and a substantial reduction in winter rapeseed yield (Brown et al., 2019). Recent evidence suggests that vernalization occurs in early autumn in the field, driven not only by prolonged cold but also by the absence of warmth (Hepworth et al., 2018; O’Neill et al., 2019).

Growth temperature emerges as a crucial factor influencing rapeseed yield, seed oil content, and fatty acid composition (Deng & Scarth, 1998; Marjanovic-Jeromela et al., 2019). Triacylglycerols (TAGs) constitute 94.1-99.1% of the total oil content in *B. napus* seeds, with oleic acid (C18:1), a monounsaturated 18-carbon fatty acid, being pivotal for oil quality and health (Shahidi, 2005; Dar et al., 2017). Breeding strategies target the development of high oleic acid varieties (Maher et al., 2007). While the impact of temperature on seed yield and quality is well-established, studies addressing the integration of temperature cues at the molecular level in developing rapeseed seeds, especially the role of epigenetic factors in response to abiotic stress during early development, are limited. Fatty acid desaturase 2 (FAD2) catalyzes the conversion of oleic acid (C18:1) to linoleic acid (C18:2) (Miquel & Browse, 1992). Mutants of *fad2* result in seeds with higher oleic acid content in both Arabidopsis and *B. napus* (Okuley et al., 1994; Hu et al., 2006). Four copies of *BnaFAD2* have been identified (Lee et al., 2013). Additionally, a GWAS analysis linked FATTY ACID DESATURASE 5 (FAD5) and Fatty acyl-ACP thioesterase B (FATB) to variations in fatty acid composition in winter rapeseed (Gacek et al., 2017).

Few studies have explored the relationship between vernalization and membrane fatty acid composition in seedlings (Thomson & Zalik, 1973; Dubert et al., 1992; Skoczowski & Filek, 1994). Winter wheat embryos exposed to different vernalization-devernalization cycles exhibited altered fatty acid composition in germinated seedlings, correlating with changes in the timing of floral induction (Dubert et al., 1992). However, the impact of a vernalization cycle temporarily interrupted by warm temperatures (>15 °C) during early growth on the fatty acid composition of seeds remains unknown.

In this study, we set up a series of experimental models to simulate shortened winters that include cold temperatures (4 °C), interrupted by one-week-long period of warmer temperature (22 °C) under laboratory conditions. These simulations, referred to as vernalization models, were implemented to investigate their impact on seed oil content and fatty acid composition in both winter and spring varieties of rapeseed. Additionally, we assessed the expression levels of three *BnaFLC* genes (*BnaFLCA02*, *BnaFLCA10*, and *BnaFLCC02*) in seedlings under these vernalization regimes. Moreover, we examined the expression patterns of various genes associated with fatty acid synthesis in both seedlings and developing seeds. The overarching goal of this study was to assess the correlations between vernalization, flowering time and several yield-related traits, and the oil content and fatty acid composition of seeds of rapeseed.

## 2. Materials and Methods

### 2.1 Plant material, seed sowing and germination

The seeds of *Brassica napus* winter varieties, Darmor and Bristol, and spring variety, Helios were kindly provided by the Germplasm Resource Information Network (GRIN, Czech Republic). Darmor has been used as the reference variety for the whole genome sequencing of *B. napus* (Chalhoub et al., 2014). Both Bristol and Helios were grown in Central Anatolia, and their suitability for cultivation in Türkiye has been confirmed (Başalma, 2004; Öztürk et al., 2008). Darmor and Bristol have been designated as late and early flowering, respectively (https://www.grin-global.org).

The experiments were conducted for three consecutive years between 2021-2023. Two seeds were sown per pot (dimensions 11×11×18 cm) filled with a soil mixture of peat and perlite (1:1) moistened with the Hoagland solution (Hoagland and Arnon, 1950). The pots were placed in a growth room with a 12-hour day/night cycle, at 22 °C, 50-55% humidity, and a light intensity of 150 μmol m^−2^ s^−1^. Plants were irrigated with fifty milliliters of Hoagland solution or distilled water at two-three day intervals to maintain soil moisture.

### 2.2 The vernalization treatment

The germinated seedlings that reached the four-five leaf stage (BBCH scale 14-15) were moved to a vernalization chamber (Nüve TK 252) at 4 °C; short days. The nonvernalized (NV) samples were left in the growth room at 22 °C; short days. The different vernalization models are illustrated in Fig. 1. At least twenty plants from each variety were subjected to vernalization for three weeks (3w); four weeks (4w); six weeks (6w); two weeks vernalization, one week devernalization at 22 °C followed by four weeks vernalization (2+4w); 8 weeks (8w) and 2 weeks vernalization, one week devernalization at 22 °C followed by 6 weeks vernalization (2+6w). For devernalization, plants were transferred to the growth room at 22 °C; short days. Previous studies have shown that the expression levels of *BnaFLC* genes in winter rapeseed varieties cannot be detected after 8 weeks of vernalization (Schiessl et al., 2019).

**Fig. 1.**
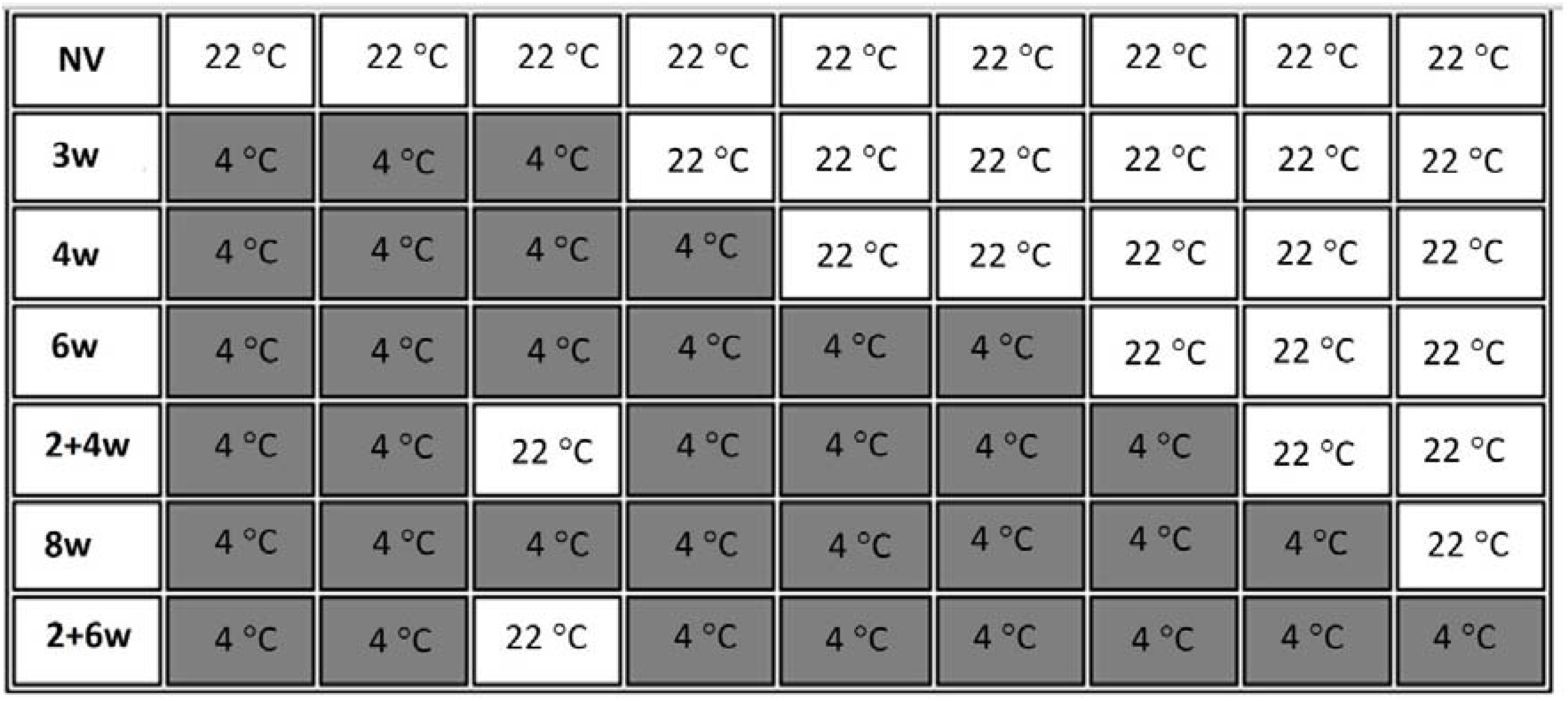
Vernalization model chart. Each box represents one week. Vernalization and devernalization took place at 4 °C and 22 °C, respectively. NV; nonvernalized, 3w; three weeks-vernalization, 4w; four weeks-vernalization, 6w; six weeks-vernalization, 2+4w; two weeks-vernalization, one week-devernalization, four weeks-vernalization, 8w; eight weeks-vernalization, 2+6w; two weeks-vernalization, one week-devernalization, six weeks-vernalization.

### 2.3 Plant growth conditions post-vernalization

Plants were transplanted to soil (slightly alkaline, clay loam textured and fertilized with diammonium phosphate before planting) outdoors on April 26, 2021, and on April 25, 2022 when the night temperature reached 10 °C and above and to a semi-controlled greenhouse with indoor heating but no cooling system on January 11, 2023. 150 W LED lamps (Philips Xitanium) provided supplementary lighting when required in the greenhouse. On October 11, 2022, Helios and Bristol seeds were directly sown in soil as winter planting. The highest and lowest daily temperatures were recorded and are given in Fig. 2A-D. The soil was irrigated every two to three days. Flowering time was recorded as the number of days after the day of transplantation when the first flower appeared or days after sowing in 2023 field plants. Yield associated physiological parameters including plant height (cm), the number of lateral branches (n), the number of pods on the main stem (n), the number of seeds per pod (n), and thousand seed weight (g) were measured.

**Fig. 2.**
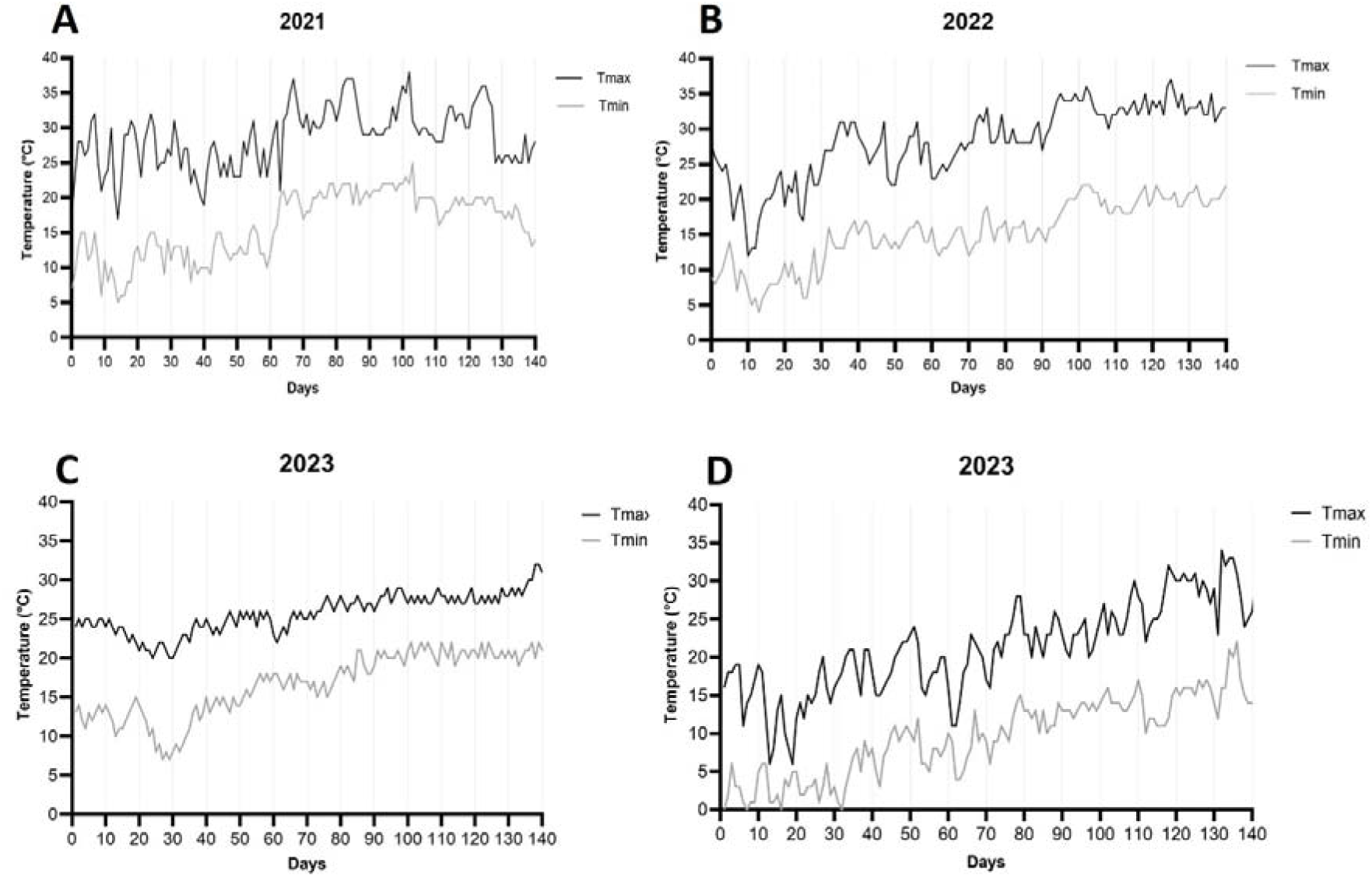
Daily maximum (Tmax) and minimum (Tmin) temperature data for Meram, Konya, Türkiye for the years 2021-2023. Tmax and Tmin starting on (Day 0) **A.** April 26, 2021; **B.** April 25, 2022; **C.** January 11, 2023 in the greenhouse; **D.** March 1, 2023 in the field.

### 2.4 RNA extraction and analysis

Leaf tissue was collected from at least three plants per vernalization model after the vernalization was completed and immediately frozen in liquid nitrogen. Siliques on the main stem that reached BBCH growth scale 80 (i.e. beginning of ripening, seeds green, filling pod cavity) were collected, developing seeds were separated and immediately frozen in liquid nitrogen. Tissue was homogenized and RNA extraction was performed as described in Queseda et al. (2003). Total RNA was DNase treated with the Turbo DNA-free kit (Thermo Fisher Scientific). For quantitative real-time RT-PCR, one μg total RNA was reverse-transcribed with the iScript cDNA synthesis kit (Bio-Rad) according to the manufacturer’s instructions. Six μL cDNA (diluted 1:5 with distilled water) was used per qRT-PCR reaction in a 15-μL reaction volume using SYBR Green (BIO-RAD, iTaq Universal SYBR Green Supermix) and a BIO-RAD CFX96 real time PCR instrument. Expression was normalized to Ubiquitin using the comparative cT method.

Primers for *B. napus* genes including *BnaFAD2.A5* and *BnaFAD2.C5*, *BnaFAD5* (*BnaA05g23670D*), *BnaFATB* (*BnaA05g23790D*), *BnaMCOA* (*AAE13*; BnaA05g23520D), *BnaWD40* (*BnaA05g24090*), *BnaFLCA02*, *BnaFLCA10*, *BnaFLCC02*, and *Ubiquitin* as internal control were either designed using the Primer3Plus program (http://www.bioinformatics.nl/cgibin/primer3plus/primer3plus.cgi) or according to the relevant reference indicated in Supplementary Table S1. The gene sequences are available at http://www.genoscope.cns.fr/brassicanapus/.

### 2.5 Total lipid extraction and gravimetric lipid determination

Seeds were harvested from at least three plants per vernalization model and dried. Total lipid extraction was done as described by Li et al. (2006). Briefly, two mL of isopropanol was added to 100 mg seeds in a glass tube with a Teflon© cap. The tubes were incubated at 85 °C in a water bath for 10 minutes. The seeds were then homogenized using a glass-glass homogenizer. Three mL of hexane was added to the sample and vortexed. Following a five-minute incubation at room temperature, 2.5 mL of 15% w/v sodium sulfate was added, leading to phase separation. The upper phase was transferred to a new glass tube and evaporated under nitrogen. The total oil content (%) was calculated gravimetrically.

### 2.6 Determination of fatty acid composition of seeds

The fatty acid composition of *B. napus* seeds was determined by gas chromatography (GC). The lipid extracts were subjected to transesterification with 0.5 mL of 2 N potassium hydroxide as catalyst. 10 mL of n-hexane was added to the samples and mixed. The mixture was centrifuged at 5000 rpm for 15 minutes, and the upper phase was collected for GC analysis using SHIMADZU (GCMS-QP2020) instrument with a column (RESTEK; Rxi-5 Sil MS 30 Meter 0.25 mm ID 0.25 μm df). The column was operated at 250 °C and 34.17 psi (split flow: 207.1 mL/min, total flow: 215.9 mL/min). Fatty acid methyl esters were quantified as percentages by comparing peak areas with standards.

### 2.7 Statistical analysis

All experiments were carried out with at least three biological replicates and results are expressed as mean ± standard error (SE). The mean values of data were analyzed by one-way analysis of variance (ANOVA) and unpaired t-tests (Mann-Whitney U test) was employed for group comparisons. Statistical significance was determined at the 5% level (*p* < 0.05). Correlations among different parameters were calculated using Pearson’s correlation coefficient (*r*). The statistical analyses were performed using GraphPad Prism 9 software.

### 3. Results

### 3.1 Expression profiles of *BnaFLC* genes and flowering time under different vernalization models

We addressed the effect of the vernalization models (NV, 3w, 4w, 6w, 2+4w, 8w and 2+6w) on the expression levels of three *BnaFLC* genes; *BnaFLCA02*, *BnaFLCA10* and *BnaFLCC02* as well as the flowering time in rapeseed winter varieties Darmor and Bristol and spring variety Helios. Initially, the expression levels of *BnaFLC* genes in the nonvernalized (NV) leaf samples of the three varieties were within a comparable range (Fig. 3). Notably, after three weeks of vernalization (3w), a significant disparity in the downregulation rates of *FLC* genes was observed in Darmor, Bristol, and Helios. Darmor, characterized as a late-flowering winter variety, exhibited only a marginal reduction in *BnaFLCA02* and *BnaFLCC02* expression levels and a three-fold reduction in *BnaFLCA10* expression (Fig. 3A-C). In contrast, Bristol, an early-flowering winter variety, displayed a higher reduction rate in all three *BnaFLC* genes compared to Darmor, yet not as pronounced as the sharp decrease observed in Helios (Fig. 3). Specifically, and *BnaFLCC02* levels in Helios were reduced approximately five-fold, whereas *BnaFLCA10* showed an eleven-fold reduction after three weeks of vernalization (3w) (Fig. 3G-I). All three *FLC* genes were undetected in Helios after six weeks of vernalization (6w). Despite being significantly lower than the nonvernalized (NV) state, Darmor and Bristol still displayed expression of *BnaFLCA02*, *BnaFLCA10*, and *BnaFLCC02* after eight weeks of vernalization (8w). *BnaFLCA10* emerged as the most responsive to vernalization in all three varieties. Notably, Darmor’s *BnaFLCC02* expression level was slightly higher in plants exposed to eight-week vernalization with a one-week interruption (2+6w) compared to those exposed to eight-week vernalization without any interruption (8w) (Fig. 3C). Conversely, in Bristol, 2+6w *BnaFLCC02* expression was higher than 8w expression (Fig. 3F).

**Fig. 3.**
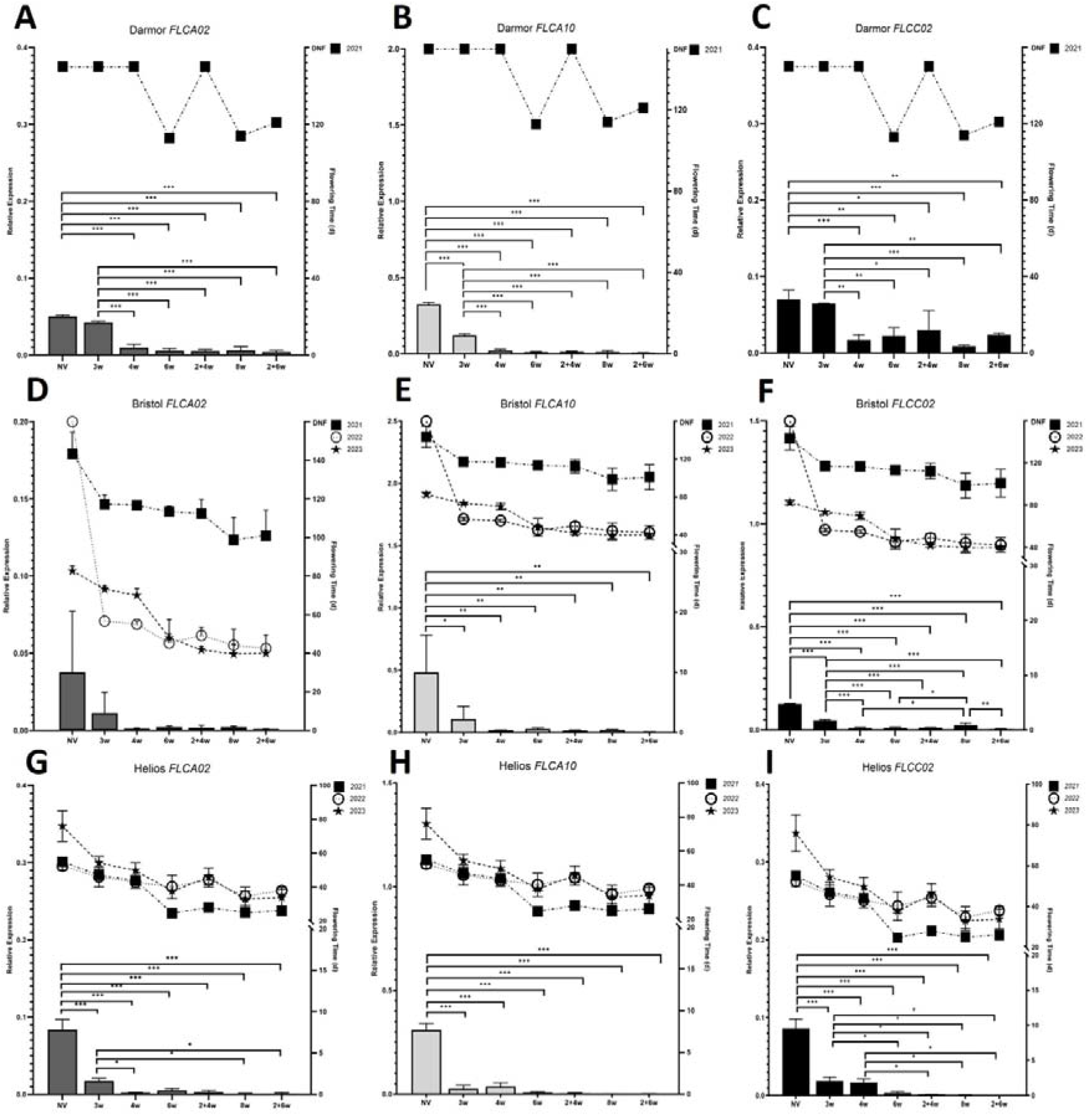
*BnaFLC* expression and flowering times of rapeseed plants under the vernalization models. **A.** *BnaFLCA02*; **B.** *BnaFLCA10*; **C.** *BnaFLCC02* relative expression levels in Darmor, **D.** *BnaFLCA02*; **E.** *BnaFLCA10*; **F.** *BnaFLCC02* relative expression levels in Bristol, **G.** *BnaFLCA02*; **H.** *BnaFLCA10*; **I.** *BnaFLCC02* relative expression levels in Helios. Histograms show mean values ± SEM for two independent PCR amplifications on three biological replicates. The left panel y axis shows the relative expression after normalization to *Ubiquitin* expression. The right panel y axis shows the flowering time as the days after transplantation when the first flower was seen. DNF: did not flower. * *p* < 0.05; ** *p* < 0.01; *** *p* < 0.001

In the initial two years (2021 and 2022) of the study, seedlings were transplanted into soil and grown under field conditions post-vernalization. Bristol and Helios were previously grown in Central Anatolia, while there is no information available about the cultivation of Darmor in Türkiye (Başalma, 2004; Öztürk et al., 2008). Darmor exhibited severe growth defects and extreme sensitivity to pests in 2021, with only a few plants from the 6w, 8w, and 2+6w vernalization models flowering. Plants from other vernalization models (NV, 3w, 4w, 2+4w) did not flower 160 days after transplantation (Fig. 3A-C). Consequently, we determined that Darmor is unsuitable for planting in the soil and weather conditions of Konya, Türkiye, and discontinued its cultivation in the following years of the study.

In 2021 and 2023, plants from all vernalization models of Bristol flowered. However, in 2022, Bristol NV plants did not flower, while the rest flowered significantly earlier than in the 2021 field season. Although flowering was observed, plants from 3w and 4w vernalization models failed to complete their growth cycle in 2022. Across all three years, there was no statistically significant difference in flowering times between 6w and 2+4w or 8w and 2+6w vernalization models in Bristol (Fig. 3D-F). Helios NV plants consistently flowered earlier than Bristol NV plants over the three years (Fig. 3G-I).

Throughout the 2021 and 2022 field seasons, flowering times of plants in all vernalization models of Helios were similar. In 2023, under greenhouse conditions, while flowering times were longer compared to previous years, a faster rate of reduction in flowering time between NV and 3w vernalization models was observed than in 2021 and 2022. A slight increase in mean flowering times of plants in the 2+4w vernalization model compared to 6w was noted across all three years (∼3-4 days in 2021 and 2022; ∼8 days in 2023). There was no meaningful difference between 2+6w and 8w in 2021 and 2023, with a slight increase in 2+6w compared to 8w (∼5 days) in 2022. These results were not statistically significant, possibly due to the variation among flowering times of plants and the small sample size (3≤n<10).

During the 2022-2023 growing season, alongside transplanting the seedlings in the greenhouse, Bristol and Helios seeds were directly sown in soil in October 2022. The plants from the Bristol and Helios varieties flowered in early April 2023; 179±2 and 182±4 days after sowing, respectively.

We analyzed the relationship between vernalization, represented by vernalization models NV, 3w, 4w, 6w and 8w expressed as days of vernalization, and the expression levels of the *BnaFLC* genes, as well as the annual flowering time. The Pearson correlation coefficients (*r*) between each pair of variables are presented in Fig. 6. As anticipated, a robust negative correlation between vernalization and flowering times was observed consistently across all three years for both Bristol and Helios varieties (*p* < 0.05). The expression levels of *BnaFLCA02*, *BnaFLCA10*, and *BnaFLCC02* of both Bristol and Helios varieties exhibited a negative correlation with vernalization and a positive correlation with flowering time consistent with the role of FLC in vernalization response.

### 3.2 Yield related physiological traits of plants under different vernalization models

We performed a comprehensive analysis of various yield-related traits in rapeseed plants, including parameters such as plant height, number of lateral branches, number of pods on the main stem, number of seeds per pod, and thousand seed weight. This investigation spanned three consecutive years and involved the evaluation of Bristol and Helios varieties under different vernalization models.

In 2023, the plant height of both Bristol and Helios varieties cultivated in the greenhouse reached its maximum, surpassing the heights of field-grown plants in both 2021 and 2022. (Fig. S1A-B). Intriguingly, the 2+6w vernalization model resulted in the shortest plants during the same year (*p* < 0.05). Conversely, during the 2021 planting season, the tallest plants were observed in the Bristol variety under the 2+6w vernalization model (*p* < 0.05). Notably, a substantial variability in physiological traits was discerned across the three years, underscoring the influence of multifactorial events, possibly attributable to specific growth conditions prevailing in each year. An exception to this trend was noted in the number of lateral branches in the Helios variety, which exhibited a subtle yet consistent decline with increasing vernalization time, irrespective of the planting year (Fig. S1D).

In the 2023 planting season, the vernalization models 2+4w and 2+6w in both Bristol and Helios varieties yielded plants with a reduced number of pods on the main stem compared to those subjected to 6w and 8w vernalization, respectively, in greenhouse conditions. This observation suggests an interplay between vernalization conditions and pod development.

We calculated the Pearson correlation coefficients (*r*) to gain insight into the interplay among vernalization, flowering time, and yield-related physiological traits (Fig. 6). In 2021, there was a positive correlation between the number of pods on the main stem and a negative correlation between the seed number per pod and the duration of vernalization in Bristol (*p* < 0.05) (Fig. 6A). In 2022, Bristol NV, 3w, and 4w plants initiated flowering but failed to progress into fully developed, healthy plants. Consequently, our analysis for this specific year excluded physiological traits other than plant height (Fig. 6B). In 2023, although a positive correlation was noted between the number of pods on the main stem and the vernalization model in Bristol, the result was not statistically significant (Fig. 6C). Turning to Helios, a robust positive correlation between thousand seed weight and the days of vernalization (i.e., vernalization model) was evident in 2022 (*p* < 0.001), but this association was not observed in other years (Fig. 6E). On the contrary, the number of lateral branches displayed a robust positive correlation with flowering time in both 2021 and 2022, and a subtle positive correlation in 2023 (Fig. 6D-F). These findings suggest that multiple variables, beyond vernalization and flowering time, contribute to the variability in yield-related physiological traits.

### 3.3 Effects of vernalization models on seed oil and oleic acid contents in rapeseed

We assessed the impact of vernalization models—specifically, variations in duration and the presence or absence of an interruption during vernalization—on the seed oil content and fatty acid composition of rapeseed varieties Bristol and Helios (Fig. 4).

**Fig. 4.**
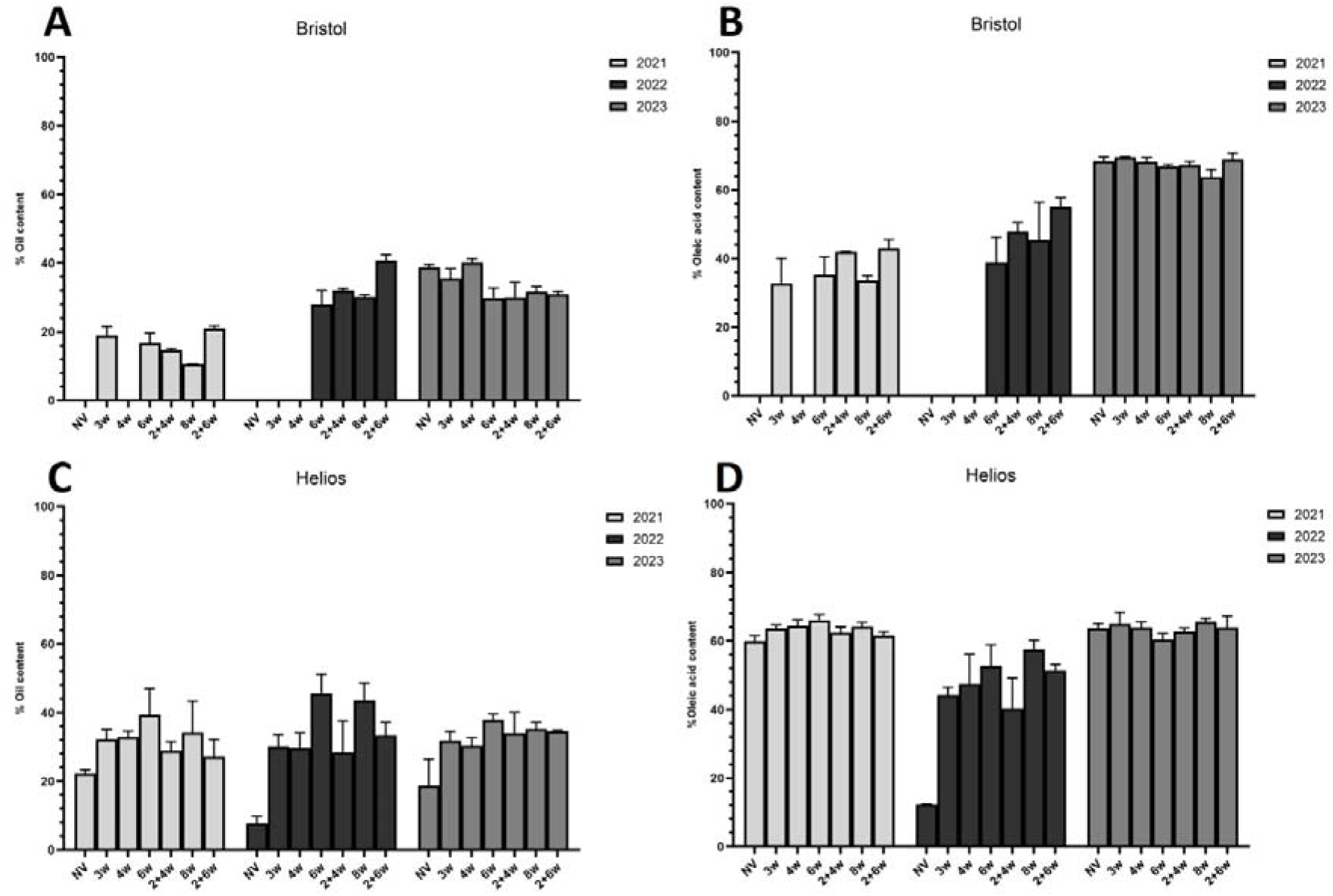
Seed oil content (% w/w) and % oleic acid content of **A** and **B** Bristol; **C** and **D** Helios under the vernalization models between 2021-2023. Histograms show mean values ± SEM for three biological replicates.

In Bristol, during both 2021 and 2022, the 2+6w vernalization model exhibited the highest oil content (% w/w) at 21% and 41%, respectively. Notably, during the same two years, a significant difference in seed oil content was observed between the 8w and 2+6w vernalization models (*p* < 0.05), underscoring the influence of vernalization interruption by warm temperatures on seed oil content. This pattern was similarly reflected in the percentage of oleic acid content (Fig. 4B). Across all three years, seeds from plants vernalized for 2+6w consistently demonstrated higher oleic acid content compared to those vernalized for 8w.

In 2023, the 4w vernalization model exhibited the highest oil content at 40% (Fig. 4A). There was no significant difference in seed oil content (% w/w) between the 8w and 2+6w vernalization models in 2023. The relationship between seed oil content (%) and flowering time was notably strong in 2021 (*r* = 0.999; *p* < 0.05) and 2023 (*r* = 0.839; *p* > 0.05) (see Fig. 6A and C). Furthermore, a robust positive correlation between flowering time and percentage of oleic acid content was observed in 2023 (*r* = 0.894; *p* < 0.05).

In Helios, seeds from non-vernalized (NV) plants exhibited the lowest oil content across all three years. In 2022, NV plants were adversely affected by the weather conditions and pests, leading to the production of small, unhealthy-looking seeds and a consequently, a low seed oil yield (8%). Nevertheless, even in the other two years, seeds from NV plants displayed lower oil content than those of the other vernalization models (Fig. 4C).

6w plants had the highest seed oil contents across all three years, surpassing those of 2+4w plants. Similarly, the seed oil contents of 8w plants in 2021 and 2022 exceeded those of 2+6w plants (*p* < 0.05), though the difference was not statistically significant in 2023. Oleic acid contents (%) showed significant differences between the 6w and 2+4w, as well as the 8w and 2+6w vernalization models in 2022. A similar trend was evident in 2021 and 2023, albeit with more subtle differences.

We calculated the relative amounts of saturated (SFA), monounsaturated (MUFA), and polyunsaturated (PUFA) fatty acids for each vernalization model throughout the 2021-2023 period (refer to the supplementary file). Remarkably, there was a significant increase in MUFA during interrupted vernalization models (2+4w and 2+6w) compared to uninterrupted vernalization (6w and 8w, respectively) across all three years in the Bristol variety. A similar effect was observed in Helios during the 2021 and 2022 field seasons, albeit with a more subtle impact.

The relationship between vernalization duration, flowering time, and seed oil content in Helios remained robust across all field seasons (Fig. 6D-F). The percentage of seed oil content exhibited a positive correlation with increasing vernalization duration in 2021 (*r* = 0.825; *p* > 0.05), 2022 (*r* = 0.944; *p* < 0.05), and 2023 (*r* = 0.884; *p* < 0.05). Oleic acid content (%) also displayed a positive correlation with vernalization duration in 2021 (*r* = 0.790; *p* > 0.05) and 2023 (*r* = 0.921; *p* < 0.05). This interplay between oil content and vernalization was further highlighted by the strong negative correlation between gene expression levels of *BnaFLCA02*, *BnaFLCA10*, and *BnaFLCC02* and seed oil content in Helios across all three years (Fig. 6D-F). In summary, our findings underscore the profound impact of vernalization on seed oil content and fatty acid composition in rapeseed.

### 3.4 The expression profiles of the fatty acid synthesis-related genes in leaf tissues across the vernalization models

We investigated the expression profiles of several genes related to fatty acid synthesis, including *BnaFAD2*, *BnaFAD5*, *BnaFATB*, *BnaMCOA* (*AAE13*), and *BnaWD40*, in the leaf tissue of Darmor, Bristol, and Helios varieties under various vernalization models. In NV samples, the expression levels of all five genes were higher in Helios compared to Darmor and Bristol (Fig. S3). Notably, there was no uniform response to vernalization in gene expression levels among different varieties.

For instance, in Darmor, *BnaFAD2*, responsible for catalyzing the conversion of oleic acid (C18:1) to linoleic acid (C18:2), exhibited an increase in expression upon cold exposure until six weeks (6w and 2+4w) of vernalization, after which a sudden drop occurred at 8w. Conversely, in Helios, there was an abrupt decline in *BnaFAD2* levels after three weeks (3w) of cold exposure, which persisted after 6w and 8w. Interestingly, interrupting the cold exposure after 2w with a week of warm temperature, followed by an additional four or six weeks of cold, caused a further decrease in expression levels of *BnaFAD2* (Fig. S3A).

The relative increase in expression levels of *BnaFAD5*, *BnaFATB*, *BnaMCOA*, and *BnaWD40* between the 6w vs 2+4w and 8w vs 2+6w vernalization models was evident in Helios (Fig. S3B-E). In Darmor, *BnaFAD5* expression was suppressed throughout all vernalization models (Fig. S3B). Similarly, in Bristol, the expression of *BnaFATB*, which encodes Fatty acyl-ACP thioesterase B, remained consistently low under cold conditions compared to NV (Fig. S3C). Notably, *BnaMCOA/AAE13*, encoding a malonyl-CoA synthetase, showed minimal expression in Darmor and Bristol NV samples, with its expression increasing upon cold exposure in Bristol (Fig. S3D).

The candidate gene *BnaWD40* (BnaA05g24090), encoding a protein from the Transducin/WD40 repeat-like superfamily, is implicated in the regulation of oleic acid (Gacek et al., 2017). *BnaWD40* was downregulated upon exposure to cold in Darmor at three weeks (3w), Bristol at four weeks (4w), and Helios at six weeks (6w), as illustrated in Fig. S3E.

Overall, *BnaFAD2*, *BnaFAD5*, *BnaFATB*, *BnaMCOA*, and *BnaWD40* were differentially regulated upon cold exposure in rapeseed varieties.

### 3.5 The expression profiles of the fatty acid synthesis-related genes in developing seeds across the vernalization models

We studied the expression levels of *BnaFAD2*, *BnaFAD5*, *BnaFATB*, *BnaMCOA*, and *BnaWD40* in developing seeds annually between 2021 and 2023, assessing their potential correlation with seed oil content and fatty acid composition across different vernalization models in Bristol and Helios. As mentioned earlier, Darmor plants exhibited suboptimal growth under field conditions in this study, preventing the examination of gene expression profiles in developing seeds of this variety.

The relative expression patterns of *BnaFAD2*, *BnaFAD5*, *BnaFATB*, *BnaMCOA*, and *BnaWD40* in developing seeds across all three years displayed high variability in both Bristol and Helios (Fig. 5). This stochasticity in gene expression was further evident in the statistical correlation analysis. For instance, in 2021, a negative correlation between *BnaFATB* expression and vernalization duration was observed in Bristol (*r* = −0.997; *p* < 0.05) (Fig. 5A). However, in 2023, a positive correlation was identified (*r* = 0.776; *p* > 0.05) (Fig. 5C). Similarly, in 2021, Bristol *BnaFAD2* expression was positively correlated with *BnaFLCA02* (*r* = 1.000; *p* < 0.01), *BnaFLCA10* (*r* = 0.997; *p* < 0.05), and *BnaFLCC02* (*r* = 0.940; *p* > 0.05), whereas this correlation was not observed in 2023 (see Fig. 5A and C).

**Fig. 5.**
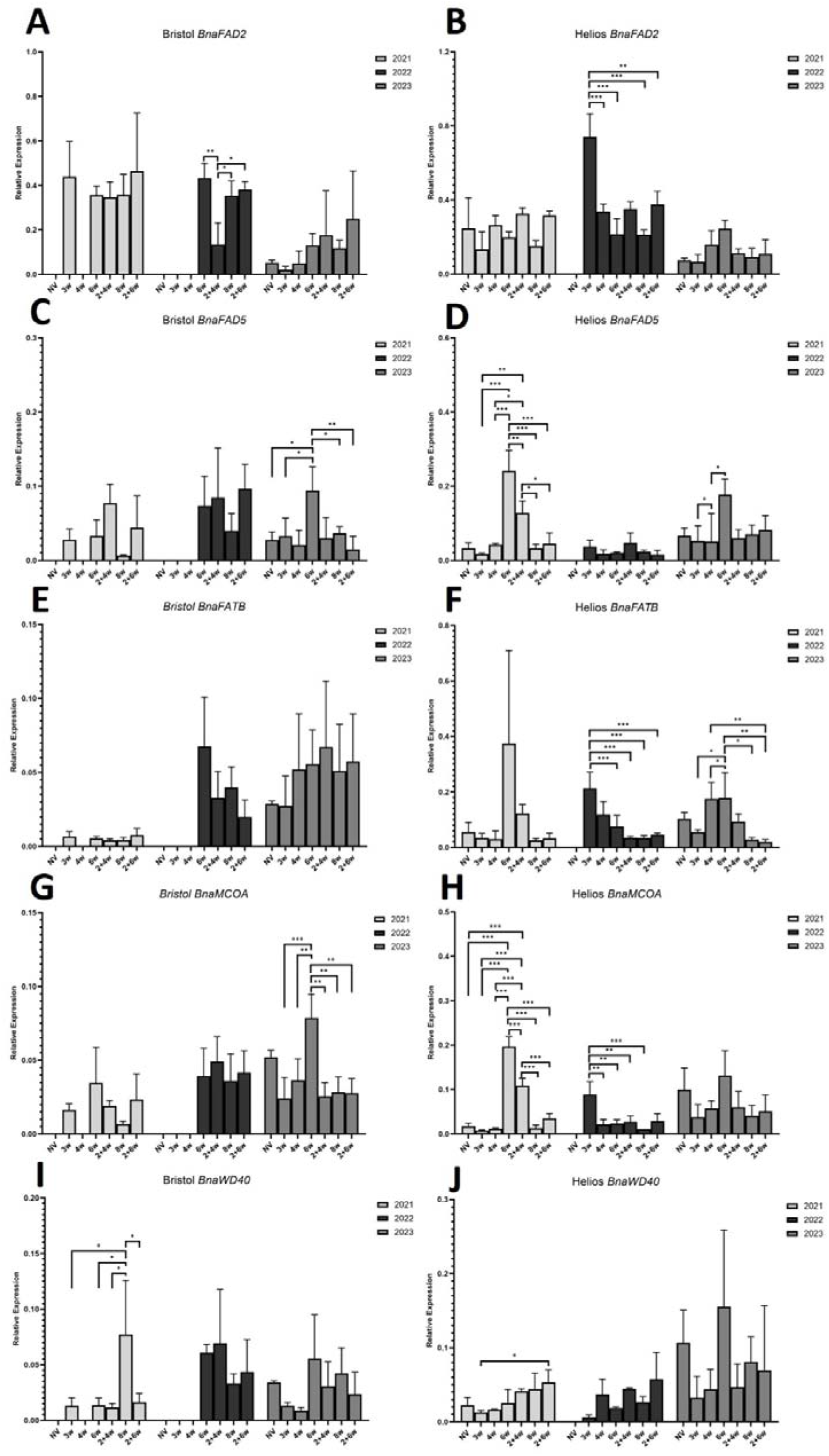
The expression profiles of the fatty acid synthesis-related genes in developing seeds under the vernalization models between 2021-2013. Relative expression levels of *BnaFAD2* in **A.** Bristol, **B.** Helios; *BnaFAD5* in **C.** Bristol, **D.** Helios; *BnaFATB* in **E.** Bristol, **F.** Helios; *BnaMCOA* in **G.** Bristol, **H.** Helios; *BnaWD40* in **I.** Bristol, **J.** Helios are shown as histograms of mean values ± SEM for three biological replicates. * *p* < 0.05; ** *p* < 0.01; *** *p* < 0.001

**Fig. 6.**
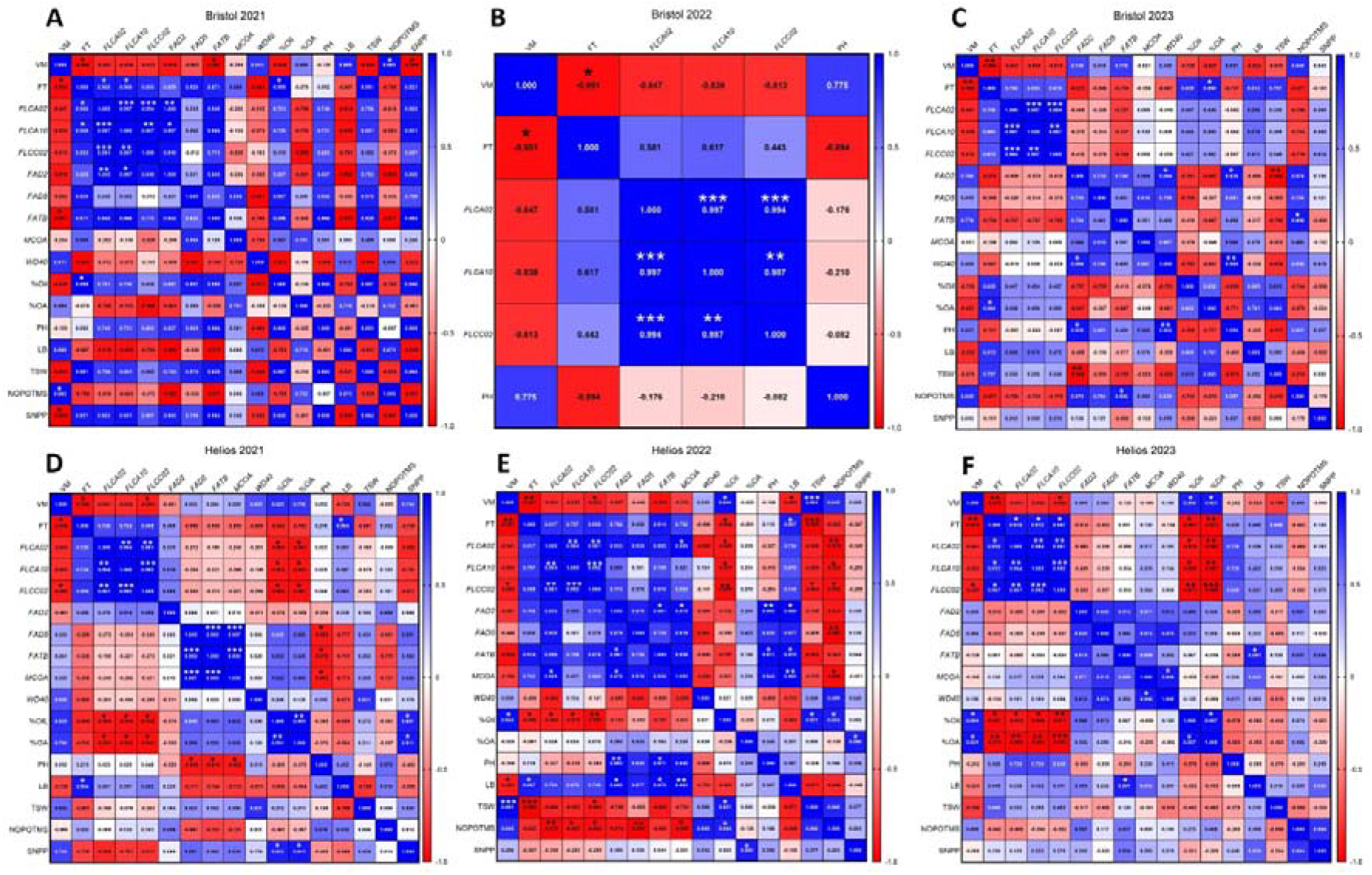
Heat maps of Pearson correlation coefficients (−1 ≤ *r* ≤ 1) between the vernalization models (NV, 3w, 4w, 6w and 8w) and the flowering time, *BnaFLC* gene expressions, fatty acid-related gene expressions, seed oil content, oleic acid content and the yield-related traits of **A.** Bristol in 2021; **B.** Bristol in 2022; **C.** Bristol in 2023; **D.** Helios in 2021; **E.** Helios in 2022; **F.** Helios in 2023. VM: vernalization model, FT: flowering time, PH: plant height, LB: lateral branches, TSW: thousand seed weight, NOPOTMS: number of pods on the main stem, SNPP: seed number per pod. * *p* < 0.05; ** *p* < 0.01; *** *p* < 0.001

The negative correlation between Bristol *BnaFAD2* expression and oleic acid content (%) in 2021 (*r* = - 0.801; *p* > 0.05) and 2023 (*r* = −0.847; *p* > 0.05), was not observed in Helios (see Fig. 5).

In summary, our findings indicate the absence of consistent correlations between the expression patterns of fatty acid-related genes that were included in this study in developing rapeseed seeds and vernalization, flowering time, oil content, and the oleic acid content.

## 4. Discussion

Vernalization is a drive for a major developmental switch in the life cycle of many plants. In this study, we aimed to elucidate the relationship between varying durations of vernalization and vernalization interrupted by warm temperature intervals and the seed oil content and fatty acid composition in winter and spring varieties of rapeseed. For this purpose, we established an experimental set-up in which four-five leaf stage rapeseed seedlings were exposed to vernalization for different durations, which we collectively name as ‘vernalization models’. As an indicator of the effectiveness of vernalization and to measure the ‘vernalization response’ of each rapeseed variety, we assessed the expression levels of three *BnaFLC* genes—namely, *BnaFLCA02*, *BnaFLCA10*, and *BnaFLCC02*—under the various vernalization models. The significance of these three paralogs in determining flowering time and vernalization requirements across different crop types has been well-documented in previous studies (Hou et al., 2012; Raman et al., 2016; Schiessl et al., 2019; Tudor et al., 2020; Yin et al., 2020).

Our findings, based on the three varieties investigated in this study, indicate that the rate at which the expression levels of the three *BnaFLC* paralogs decline (i.e., the vernalization response) is a more pertinent indicator of crop-type than the initial (NV) levels of *FLC* genes. Notably, the NV levels of *BnaFLCA02*, *BnaFLCA10*, and *BnaFLCC02* in all three varieties did not exhibit a significant difference, contrasting with findings from previous studies (Schiessel 2019; Tudor et al., 2020). It is plausible that the Helios variety, particularly among spring-types, stands as an exception with initially high *BnaFLCA10* levels (Schiessel et al., 2019). Additionally, it remains possible that the other six *FLC* paralogs, unexplored in this study, also contribute to the vernalization requirement of rapeseed, as demonstrated previously by Calderwood et al. (2020).

The rapeseed variety Helios has traditionally been classified as a spring-type in multiple studies (Jorgensen & Andersen, 1994; Brown et al., 1995; Öztürk et al., 2008). However, in the context of the present study, we successfully cultivated Helios as a winter-type by planting it in the field during the autumn season. This observation underscores the remarkable genetic diversity inherent in rapeseed varieties, especially concerning flowering time and vernalization requirements. The intricate nature of this diversity further complicates the unequivocal classification of accessions into discrete crop-types.

We observed a significant delay in flowering for Helios, including non-vernalized (NV) plants, during 2023 under semi-controlled greenhouse conditions when compared to the preceding two years (2021 and 2022). A plausible explanation is that even the NV plants were effectively vernalized after transplantation to soil in 2021 and 2022, but not in 2023, considering rapeseed’s demonstrated responsiveness to vernalization up to 17 °C (O’Neill et al., 2019).

It’s worth noting that while the greenhouse lacked fully automated temperature control, the daily mean temperatures exhibited less fluctuation than the outdoor conditions experienced in 2021 and 2022. As highlighted by Burghardt et al. (2016), the limited temperature fluctuations in the greenhouse may also have contributed to the delayed flowering times observed across all models of Helios in 2023. Additionally, the combined impact of high light exposure and variable temperatures could have further influenced the flowering times (Burghardt et al., 2016).

It is important to emphasize that, in contrast to Helios, Bristol plants did not exhibit delayed flowering under greenhouse conditions compared to outdoor field conditions.

All three *BnaFLC* levels in Helios plants were undetected after eight weeks of vernalization, consistent with full vernalization. Consequently, the interruption of vernalization by a one-week warm interval did not affect the flowering times of plants under the 2+6w model compared to the 8w model. However, Helios plants exhibited delayed flowering under the interrupted vernalization model (2+4w) compared to the uninterrupted model (6w) across all three growing seasons. In contrast, Bristol plants were unaffected by the interruption of vernalization, emphasizing the genotypic differences in sensitivity to vernalization between the two varieties.

In a prior study, O’Neill et al. (2019) implemented a warming plot system in the field during autumn, revealing that flowering in winter rapeseed was delayed by only five to seven days, while floral induction was delayed by at least twenty days. In the current study, despite a significantly shorter warm interval (one week vs. one month), we observed a slight delay in flowering time for the spring variety Helios, but not for Bristol. Future investigations could explore the floral transition in both varieties under a one-week interruption, and whether winter varieties like Darmor and Bristol would display a similar delay in flowering times under a warming plot experimental setup.

Furthermore, it would be intriguing to examine whether the atypical spring-type Helios also exhibits bud dormancy akin to the winter annual types studied by Lu et al. (2022). Lu et al. suggested that the bud dormancy might contribute to the relative difference in timing between floral transition and flowering times.

The influence of temperature on the seed oil content and fatty acid composition of rapeseed during seed filling has been well-established in the literature (Canvin, 1965; Trémolières et al., 1978; Deng & Scarth, 1998; Aksouh-Harradj et al., 2006; Zhou et al., 2018). The synthesis and storage of seed components predominantly occur within the first two to five weeks after flowering (Fowler & Downey, 1969; Rakow & McGregor, 1975). Consequently, any temperature variations during this critical period have significant implications for seed oil content and composition. The alignment of flowering time with optimal environmental conditions plays a crucial role in fine-tuning plant development for reproductive success. The variation in seed oil content and fatty acid composition observed across different vernalization models for Helios and Bristol can be partially attributed to the differences in flowering times.

The impact of vernalization on the lipid content of seedlings has been explored in previous studies involving wheat, rye, and rapeseed plants (Redshaw & Zalik, 1968; Thomson & Zalik, 1973). These investigations primarily sought to comprehend the structural and physiological alterations in plant cell membranes as an adaptive response to vernalization, which involves prolonged exposure to cold temperatures. While fatty acids predominantly constitute a major component of plant membranes in vegetative tissues, in seeds, they are primarily utilized for storage. Notably, to our knowledge, the specific influence of vernalization on the lipid composition of seeds has not been examined to date.

Our results highlight a correlation between vernalization and seed oil content and composition in rapeseed. Intriguingly, this relationship may be partially independent of plant flowering times, as observed in Bristol plants under 8w vs. 2+6w vernalization models. Despite not showing statistically significant variation in flowering times, Bristol plants exhibited significant differences in seed oil and oleic acid contents.

Interestingly, apart from its role in regulating the flowering pathway, FLC is associated with other developmental processes, including the temperature-dependent transition to seed germination in Arabidopsis (Chiang et al., 2009). The identification of genome-wide targets directly bound by FLC includes genes involved in the lipid metabolism pathway (Deng et al., 2011). Speculatively, the low oil content in non-vernalized (NV) Helios seeds may be attributed to higher *BnaFLC* levels directly interfering with lipid synthesis during seed development, rather than differences in flowering times.

The epigenetic silencing of *FLC* is reset during embryogenesis in Arabidopsis (Sheldon et al., 2008). It remains unknown whether all nine copies of *BnaFLC* follow a similar expression pattern throughout the life cycle of vernalized rapeseed plants. Given the genotypic diversity of *BnaFLC* alleles in various crop types, a comprehensive analysis is necessary to investigate *FLC* expression during seed development in a wide range of rapeseed accessions.

Understanding seed development as a complex mechanism involving intricate pathways implies that multiple factors may contribute to the formation of seed components. A recent study unraveled the direct role of circadian rhythm regulators in initiating fatty acid synthesis in Arabidopsis seeds (Kim et al., 2023), further underscoring the multifaceted nature of seed development.

The role of epigenetic factors in regulating fatty acid synthesis in plant seeds in response to temperature changes during early development is still an open question that requires further elucidation. While research has uncovered the involvement of epigenetic mechanisms in various plant developmental processes, including flowering time and stress responses, their specific impact on the regulation of fatty acid synthesis during seed development remains an area that needs more investigation. Understanding the interplay between genetic and epigenetic factors could contribute to a more comprehensive understanding of how plants adapt to environmental conditions during seed development, with potential implications for improving crop traits and yield.

## Conclusion

Present study explores the intricate connections among vernalization, flowering times, and seed traits in winter and spring rapeseed varieties. By employing diverse vernalization models and assessing *BnaFLC* gene expression, we have revealed key insights into the underlying mechanisms.

Our results emphasize that the rate of decline in *BnaFLC* paralog expression, indicating the vernalization response, plays a pivotal role in determining crop type. This response appears more crucial than the initial non-vernalized levels of *FLC* genes. The genetic diversity observed, particularly in the unique characteristics of the Helios spring-type, challenges the straightforward classification of rapeseed varieties.

Additionally, the lipid content and fatty acid composition in response to vernalization models suggests the potential independent influence of vernalization on seed characteristics beyond its impact on flowering times. The association between *FLC* genes and lipid metabolism pathway genes opens intriguing avenues for further research into their direct role in seed development.

This study contributes valuable insights into the complex nature of vernalization and its cascading effects on various aspects of rapeseed biology. The potential role of epigenetic factors in regulating fatty acid synthesis during seed development presents an intriguing area for future research, offering insights into plant adaptation to environmental conditions.

In summary, our comprehensive investigation provides valuable insights into the complex interactions of vernalization, gene expression, flowering times, and seed characteristics in rapeseed, laying the foundation for further exploration and refinement of our understanding in this field.

## Acknowledgements

This study was funded by TUBITAK grant number 120Z278 to Ç. Sönmez. B. E. Kıvrak was funded by TUBITAK 2209-A undergraduate fellowship program.

---

**Fig. S1.**
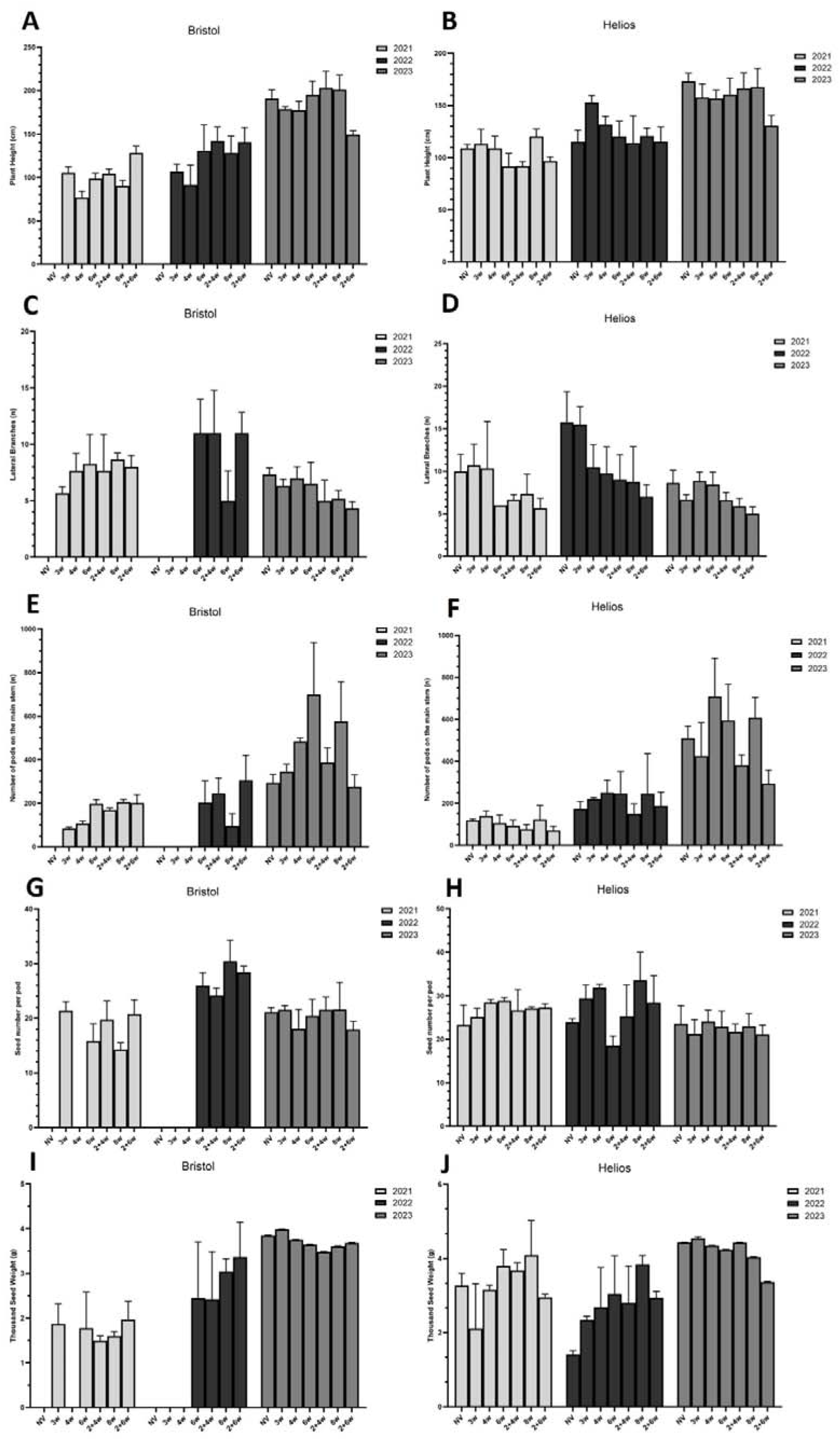
Yield-related physiological traits under the vernalization models between 2021-2023. Plant height (cm) of **A.** Bristol, **B.** Helios; lateral branches (n) of **C.** Bristol, **D.** Helios; number of pods on the main stem of **E.** Bristol, **F.** Helios; seed number per pod of **G.** Bristol, **H.** Helios; thousand seed weight (g) of **I.** Bristol, **J.** Helios. Histograms show mean values ± SEM for at least three biological replicates.

**Fig. S2.**
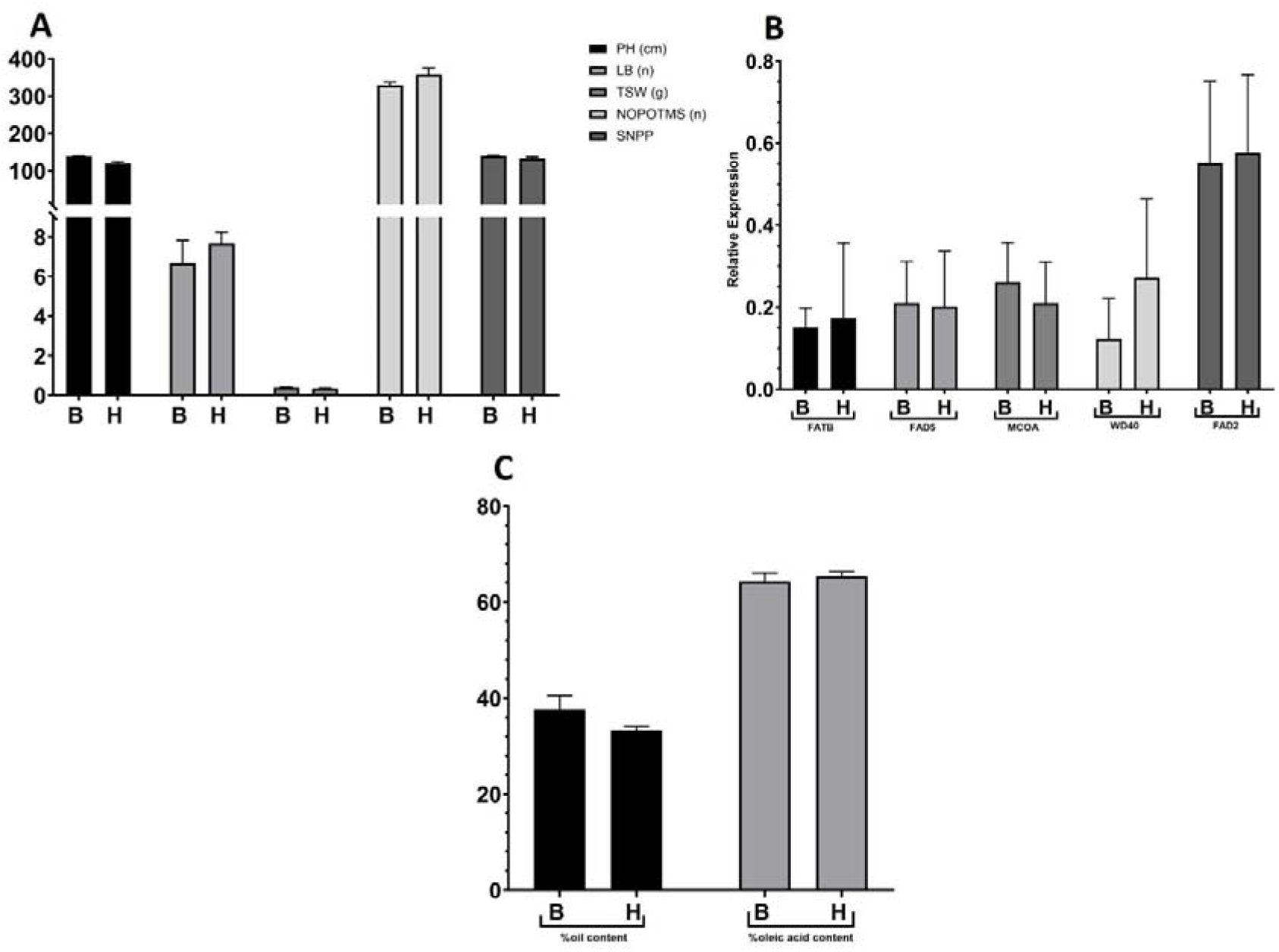
The comparison of Bristol and Helios varieties grown in the field by direct sowing in Fall 2022. **A.** Yield-related traits; plant height (PH), lateral branches (LB), thousand seed weight (TSW), number of pods on the main stem (NOPOTMS), and seed number per pod (SNPP) are shown as histograms of mean values ± SEM for at least three biological replicates. **B.** The expression profiles of the fatty acid synthesis-related genes in developing seeds, and **C.** Seed oil content (% w/w) and oleic acid content (%) are shown as histograms of mean values ± SEM for three biological replicates.

**Fig. S3.**
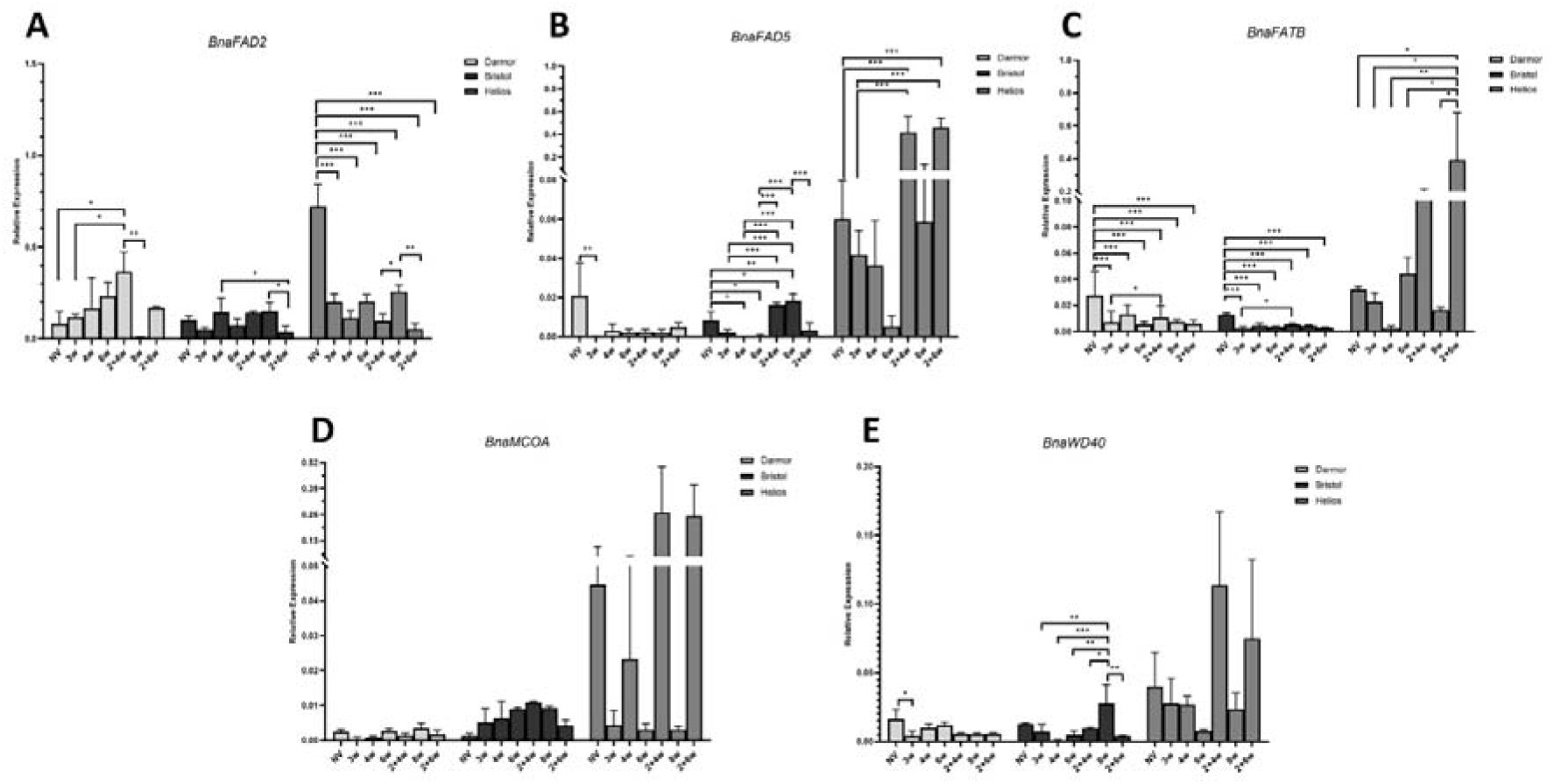
The expression profiles of the fatty acid synthesis-related genes in leaves of seedlings under the vernalization models in Darmor, Bristol and Helios. Relative expression levels of **A.** *BnaFAD2*; **B.** *BnaFAD5;* **C.** *BnaFATB*; **D.** *BnaMCOA*; **E.** *BnaWD40* are shown as histograms of mean values ± SEM for three biological replicates. * *p* < 0.05; ** *p* < 0.01; *** *p* < 0.001

## Notes

### Competing Interest Statement

The authors have declared no competing interest.

### Summary of Updates

A discussion and a conclusion section have been added. Minor typos have been corrected.

## References

Aksouh-Harradj, N. M., Campbell, L. C., & Mailer, R. J. (2006). Canola response to high and moderately high temperature stresses during seed maturation. Canadian journal of plant science, 86(4), 967–980.

Basalma D. (2004) Comparison of Winter Rapeseed (*Brassica napus* ssp. oleifera L.) Varieties in terms of yield and yield components under Ankara conditions. Journal of Agricultural Sciences, 10(2), 211–217.

Bastow, R., Mylne, J. S., Lister, C., Lippman, Z., Martienssen, R. A., & Dean, C. (2004). Vernalization requires epigenetic silencing of FLC by histone methylation. Nature, 427(6970), 164–167.

Brown, J., Erickson, D. A., Davis, J. B., & Brown, A. P. (1995). Effects of swathing on yield and quality of spring canola (Brassica napus L.) in the pacific North West. In Proceedings of the 9th International Rapeseed Congress; Cambridge, UK (Vol. 1, pp. 339–341).

Brown, J. K., Beeby, R., & Penfield, S. (2019). Yield instability of winter oilseed rape modulated by early winter temperature. Scientific reports, 9(1), 6953.

Burghardt, L. T., Runcie, D. E., Wilczek, A. M., Cooper, M. D., Roe, J. L., Welch, S. M., & Schmitt, J. (2016). Fluctuating, warm temperatures decrease the effect of a key floral repressor on flowering time in Arabidopsis thaliana. New Phytologist, 210(2), 564–576.

Calderwood, A., Lloyd, A., Hepworth, J., Tudor, E. H., Jones, D. M., Woodhouse, S., & Morris, R. J. (2021). Total FLC transcript dynamics from divergent paralogue expression explains flowering diversity in Brassica napus. New Phytologist, 229(6), 3534–3548.

Canvin, D. T. (1965). The effect of temperature on the oil content and fatty acid composition of the oils from several oil seed crops. Canadian Journal of Botany, 43(1), 63–69.

Chalhoub, B., Denoeud, F., Liu, S., Parkin, I. A., Tang, H., Wang, X., & Corréa, M. (2014). Early allopolyploid evolution in the post-Neolithic *Brassica napus* oilseed genome. Science, 345(6199), 950–953

Chiang, G. C., Barua, D., Kramer, E. M., Amasino, R. M., & Donohue, K. (2009). Major flowering time gene, FLOWERING LOCUS C, regulates seed germination in Arabidopsis thaliana. Proceedings of the National Academy of Sciences, 106(28), 11661–11666.

Clarke, J. H., & Dean, C. (1994). Mapping FRI, a locus controlling flowering time and vernalization response in Arabidopsis thaliana. Molecular and General Genetics MGG, 242(1), 81–89

Crevillen, P., Sonmez, C., Wu, Z., & Dean, C. (2013). A gene loop containing the floral repressor FLC is disrupted in the early phase of vernalization. The EMBO journal, 32(1), 140–148.

Dar, A. A., Choudhury, A. R., Kancharla, P. K., & Arumugam, N. (2017). The FAD2 gene in plants: occurrence, regulation, and role. Frontiers in plant science, 8, 1789.

De Lucia, F., Crevillen, P., Jones, A. M., Greb, T., & Dean, C. (2008). A PHD-polycomb repressive complex 2 triggers the epigenetic silencing of FLC during vernalization. Proceedings of the National Academy of Sciences, 105(44), 16831–16836.

Deng, X., & Scarth, R. (1998). Temperature effects on fatty acid composition during development of low-linolenic oilseed rape (Brassica napus L.). *Journal of the American Oil Chemists’* Society, 75(7), 759–766.

Deng, W., Ying, H., Helliwell, C. A., Taylor, J. M., Peacock, W. J., & Dennis, E. S. (2011). FLOWERING LOCUS C (FLC) regulates development pathways throughout the life cycle of Arabidopsis. Proceedings of the National Academy of Sciences, 108(16), 6680–6685.

Dubert, F., Filek, M., Marcinska, I., & Skoczowski, A. (1992). Influence of warm intervals on the effects of vernalization and the composition of phospholipid fatty acids in seedlings of winter wheat. Journal of Agronomy and Crop Science, 168(2), 133–141.

Fowler, D. B., & Downey, R. K. (1970). Lipid and morphological changes in developing rapeseed, Brassica napus. Canadian Journal of Plant Science, 50(3), 233–247.

Gacek, K., Bayer, P. E., Bartkowiak-Broda, I., Szala, L., Bocianowski, J., Edwards, D., & Batley, J. (2017). Genome-wide association study of genetic control of seed fatty acid biosynthesis in Brassica napus. Frontiers in Plant Science, 7, 2062.

Geraldo, N., Bäurle, I., Kidou, S. I., Hu, X., & Dean, C. (2009). FRIGIDA delays flowering in Arabidopsis via a cotranscriptional mechanism involving direct interaction with the nuclear cap-binding complex. Plant Physiology, 150(3), 1611–1618.

Hepworth, J., Antoniou-Kourounioti, R. L., Bloomer, R. H., Selga, C., Berggren, K., Cox, D., & Howard, M. (2018). Absence of warmth permits epigenetic memory of winter in Arabidopsis. Nature communications, 9(1), 1–8.

Hoagland, D.R., and D.I. Arnon. 1950. The water culture methods for growing plants without soil. Calif. Agric. Exp. Stn. Circular Number 347 (Revised January 1950), College of Agriculture University of California, Berkeley, CA.

Hu, X., Sullivan-Gilbert, M., Gupta, M., & Thompson, S. A. (2006). Mapping of the loci controlling oleic and linolenic acid contents and development of fad2 and fad3 allele-specific markers in canola (Brassica napus L.). Theoretical and Applied Genetics, 113, 497–507.

Hou, J., Long, Y., Raman, H., Zou, X., Wang, J., Dai, S., & Meng, J. (2012). A Tourist-like MITE insertion in the upstream region of the BnFLC. A10 gene is associated with vernalization requirement in rapeseed (Brassica napus L.). BMC plant biology, 12, 1–13.

Jorgensen, R. B., & Andersen, B. (1994). Spontaneous hybridization between oilseed rape (Brassica napus) and weedy B. campestris (Brassicaceae): a risk of growing genetically modified oilseed rape. American Journal of Botany, 1620–1626.

Kim, S. C., Edgeworth, K. N., Nusinow, D. A., & Wang, X. (2023). Circadian clock factors regulate the first condensation reaction of fatty acid synthesis in Arabidopsis. Cell Reports, 42(12).

Koornneef, M., Hanhart, C. J., & Van der Veen, J. H. (1991). A genetic and physiological analysis of late flowering mutants in Arabidopsis thaliana. Molecular and General Genetics MGG, 229(1), 57–66.

Koornneef, M., Blankestijn-de Vries, H., Hanhart, C., Soppe, W., & Peeters, T. (1994). The phenotype of some late-flowering mutants is enhanced by a locus on chromosome 5 that is not effective in the Landsberg erecta wild-type. The Plant Journal, 6(6), 911–919

Lee, K. R., Sohn, S. I., Jung, J. H., Kim, S. H., Roh, K. H., Kim, J. B., & Kim, H. U. (2013). Functional analysis and tissue-differential expression of four FAD2 genes in amphidiploid Brassica napus derived from Brassica rapa and Brassica oleracea. Gene, 531(2), 253–262.

Lesk, C., Rowhani, P., & Ramankutty, N. (2016). Influence of extreme weather disasters on global crop production. Nature, 529(7584), 84–87.

Li, Y., Beisson, F., Pollard, M., & Ohlrogge, J. (2006). Oil content of Arabidopsis seeds: the influence of seed anatomy, light and plant-to-plant variation. Phytochemistry, 67(9), 904–915.

Lu, X., O’Neill, C. M., Warner, S., Xiong, Q., Chen, X., Wells, R., & Penfield, S. (2022). Winter warming post floral initiation delays flowering via bud dormancy activation and affects yield in a winter annual crop. Proceedings of the National Academy of Sciences, 119(39), e2204355119.

Maher, L., Burton, W., Salisbury, P., Debonte, L., & Deng, X. (2007, March). High Oleic, low linolenic (HOLL) specialty canola development in Australia. In Proceedings of the 12th International Rapeseed Congress (Vol. 5, pp 22–24).

Marjanović-Jeromela, A., Terzić, S., Jankulovska, M., Zorić, M., Kondić-Špika, A., Jocković, M., & Nagl, N. (2019). Dissection of Year Related Climatic Variables and Their Effect on Winter Rapeseed (Brassica napus L.) Development and Yield. Agronomy, 9(9), 517.

Michaels, S. D., & Amasino, R. M. (2001). Loss of FLOWERING LOCUS C activity eliminates the late-flowering phenotype of FRIGIDA and autonomous pathway mutations but not responsiveness to vernalization. The Plant Cell, 13(4), 935–941.

Miquel M, Browse J (1992) Arabidopsis mutants deficient in polyunsaturated fatty-acid synthesis— biochemical and genetic-characterization. Journal of Biological Chemistry 267:1502–1509

Napp-Zinn, K. (1979). On the genetical basis of vernalization requirement in Arabidopsis thaliana (L.) Heynh. La Physiologie de la Floraison, 217–220.

Okuley, J., Lightner, J., Feldmann, K., Yadav, N., Lark, E., & Browse, J. (1994). Arabidopsis FAD2 gene encodes the enzyme that is essential for polyunsaturated lipid synthesis. The Plant Cell, 6(1), 147–158.

O’Neill, C. M., Lu, X., Calderwood, A., Tudor, E. H., Robinson, P., Wells, R., & Penfield, S. (2019). Vernalization and Floral Transition in Autumn Drive Winter Annual Life History in Oilseed Rape. Current Biology.

Öztürk, Ö., Rahim, A. D. A., & Akinerdem, F. (2008). Determination of appropriate planting time for spring rapeseed varieties under Konya conditions. Selçuk Journal of Agricultural Sciences, 22(46), 6–17.

Rakow, G., & McGregor, D. I. (1975). Oil, fatty acid and chlorophyll accumulation in developing seeds of two” linolenic acid lines” of low erucic acid rapeseed. Canadian Journal of Plant Science, 55(1), 197–203.

Raman, H., Raman, R., Coombes, N., Song, J., Prangnell, R., Bandaranayake, C., & Balasubramanian, S. (2016). Genome-wide association analyses reveal complex genetic architecture underlying natural variation for flowering time in canola. *Plant*, Cell & Environment, 39(6), 1228–1239.

Redshaw, E. S., and S. Zalik 1968. Changes in lipids of cereal seedlings during vernalization. Can. J. Biochem. 46 1093–1097

Sheldon, C. C., Hills, M. J., Lister, C., Dean, C., Dennis, E. S., & Peacock, W. J. (2008). Resetting of FLOWERING LOCUS C expression after epigenetic repression by vernalization. Proceedings of the National Academy of Sciences, 105(6), 2214–2219.

Schiessl, S., Samans, B., Hüttel, B., Reinhard, R., & Snowdon, R. J. (2014). Capturing sequence variation among flowering-time regulatory gene homologs in the allopolyploid crop species Brassica napus. Frontiers in plant science, 5, 404.

Schiessl, S. V., Quezada-Martinez, D., Tebartz, E., Snowdon, R. J., & Qian, L. (2019). The vernalisation regulator FLOWERING LOCUS C is differentially expressed in biennial and annual Brassica napus. Scientific reports, 9(1), 1–15.

Shahidi F., 2005. Bailey’s Industrial Oil and Fat Products, Volume 6, Wiley-Interscience, New Jersey, USA.

Shindo, C., Aranzana, M. J., Lister, C., Baxter, C., Nicholls, C., Nordborg, M., & Dean, C. (2005). Role of FRIGIDA and FLOWERING LOCUS C in determining variation in flowering time of Arabidopsis. Plant physiology, 138(2), 1163–1173.

Skoczowski, A., & Filek, M. (1994). Changes in fatty acids composition of subcellular fractions from hypocotyls of winter rape growing at 2 C or 20 C. Plant Science, 98(2), 127–133.

Swiezewski, S., Liu, F., Magusin, A., & Dean, C. (2009). Cold-induced silencing by long antisense transcripts of an Arabidopsis Polycomb target. Nature, 462(7274), 799–802.

Trémolières, H., Trémolières, A., & Mazliak, P. (1978). Effects of light and temperature on fatty acid desaturation during the maturation of rapeseed. Phytochemistry, 17(4), 685–687.

Thomson, L. W., & Zalik, S. (1973). Lipids in rye seedlings in relation to vernalization. Plant physiology, 52(3), 268–273.

Tudor, E. H., Jones, D. M., He, Z., Bancroft, I., Trick, M., Wells, R., & Dean, C. (2020). QTL-seq identifies BnaFT. A02 and BnaFLC. A02 as candidates for variation in vernalization requirement and response in winter oilseed rape (Brassica napus). Plant biotechnology journal, 18(12), 2466–2481.

Quesada, V., Macknight, R., Dean, C., & Simpson, G. G. (2003). Autoregulation of FCA pre-mRNA processing controls Arabidopsis flowering time. The EMBO Journal, 22(12), 3142–3152.

Whittaker, C., & Dean, C. (2017). The FLC locus: a platform for discoveries in epigenetics and adaptation. Annual Review of Cell and Developmental Biology, 33, 555–575.

Yin, S., Wan, M., Guo, C., Wang, B., Li, H., Li, G., & Wang, J. (2020). Transposon insertions within alleles of BnaFLC. A10 and BnaFLC. A2 are associated with seasonal crop type in rapeseed. Journal of Experimental Botany, 71(16), 4729–4741.

Zhou, L., Yan, T., Chen, X., Li, Z., Wu, D., Hua, S., & Jiang, L. (2018). Effect of high night temperature on storage lipids and transcriptome changes in developing seeds of oilseed rape. Journal of experimental botany, 69(7), 1721–1733.

Zou, X., Suppanz, I., Raman, H., Hou, J., Wang, J., Long, Y., & Meng, J. (2012). Comparative analysis of FLC homologues in Brassicaceae provides insight into their role in the evolution of oilseed rape.

